# SOX2 and NR2F1 coordinate the gene expression program of the early postnatal visual thalamus

**DOI:** 10.1101/2023.11.27.568801

**Authors:** Linda Serra, Anna Nordin, Mattias Jonasson, Carolina Marenco, Francesca Gullo, Sergio Ottolenghi, Federico Zambelli, Michèle Studer, Giulio Pavesi, Claudio Cantù, Silvia K. Nicolis, Sara Mercurio

## Abstract

The thalamic dorsolateral geniculate nucleus, (dLGN) receives visual input from the retina via the optic nerve, and projects to the cortical visual area, where eye-derived signals are elaborated. The transcription factors SOX2 and NR2F1 are directly involved in the differentiation of dLGN neurons, based on mouse work and patient mutations leading to vision defects. However, whether they regulate each other, or control common targets is still unclear. By RNA-seq analysis of neonatal dLGN from thalamo-specific *Sox2* and *Nr2f1* mouse mutants, we found a striking overlap of deregulated genes. Among them, VGF, a cytokine transported along thalamic-cortical axons is strongly downregulated in both mutants. CUT&RUN analysis of SOX2 binding in dLGN chromatin identified a binding pattern characteristic of the dLGN. Collectively, the SOX2 and NR2F1- coregulated genes, and cognate SOX2 binding sites, contribute as a basis to understand the gene regulatory network driving the differentiation and connectivity of thalamic neurons.

## Introduction

Inherited diseases affecting vision importantly contribute to significant disability in human populations. Most frequently, the retina and the eye are involved, but other parts of the visual axis, such as thalamic nuclei and visual cortical areas can also be affected ^1–3^. A wide variety of genes are known to be mutated in such diseases, ranging from transcription factors to cell membrane receptors, to secreted signalling molecules, to second messengers in signal transduction, etc^1,2^.

Interestingly, transcription factors expressed during early development with pleiotropic functions are found to be mutated in specific cases, which raises the issue of which genes among their targets are functionally important to cause the pathological phenotype. Studies in humans and mice show that mutations of two genes, *NR2F1* and *SOX2* cause important vision defects ^1,4,5^: *NR2F1* mutations are responsible for the Bosch-Boonstra-Schaaf optic atrophy syndrome (OMIM #615722; BBSOAS), whereas *SOX2* mutations cause micro- or anophtalmia (OMIM #206900, Microphtalmia, syndromic 3; Optic nerve hypoplasia and abnormalities of the central nervous system). Thalamo-specific conditional knockout (cKO) mouse mutants show that inactivation of *Sox2* or *Nr2f1* affects, albeit at different degrees of severity, proper differentiation of the thalamic dorsolateral geniculate nucleus (dLGN) and the visual cortex, also causing alterations of the topographic connections relaying information to and from these functional locations ^6–10^. The dLGN nucleus is crucial for conveying information from the retina to the primary visual cortex.

Mouse mutants, conditionally deleted in the dLGN for either *Sox2* or *Nr2f1* using a RORα-Cre transgene, result in hypomorphic visual thalamus postnatally, reduced thalamic projections to the primary visual cortex ^7,9^, significant abnormalities of the visual cortex, and important alterations of retino-thalamic connections ^9^. We reasoned that dysregulation of a common set of target genes of NR2F1 and SOX2 might be the underlying factor causing phenotypic similarity following their thalamic (dLGN) inactivation. If this is correct, it might point to a gene set universally shared during development of the dLGN and the visual axis, as well as common to the visual phenotype of *Nr2f1* and *Sox2* mutants.

In the present work, we obtained RNA-seq data from wild-type as well as *Nr2f1* or *Sox2* dLGN mutants and identified over 500 dysregulated genes in common. We show that most of the top downregulated genes in *Nr2f1* mutants are also significantly downregulated in *Sox2* mutants (44 out of 50); likewise, the large majority of the top downregulated genes in *Sox2* mutants are significantly downregulated also in *Nr2f1* mutants. These results suggest that many of the downregulated genes identified in this study contribute to at least some of the phenotypic alterations observed in mouse mutants. Functional enrichment analysis showed that deregulated genes are highly enriched in differentiated neuronal functions (axon guidance molecules, synaptic proteins, etc.). Deconvolution analysis of RNA-seq data, based on the comparison with single-cell RNA-seq data previously obtained on wild-type visual thalamus^11^, revealed a substantial reduction in cells with a transcriptional identity of glutamatergic neurons already at early developmental stages, preceding phenotypic defects. Moreover, we employed the Cleavage Under Targets and Release Using Nuclease (CUT&RUN; ^12^) approach to identify the *in vivo* genome-wide direct SOX2 binding sites in visual thalamic nuclei. This analysis allowed us to establish: i) the first full set of SOX2 binding sites in functionally relevant differentiated neurons, ii) the subset of targets that are transcriptionally regulated by SOX2, and iii) those that likely depend on the interplay between SOX2 and NRF1.

Finally, we also identified a general SOX/NRF binding consensus sequence characteristic of the dLGN that might be used by several other transcription factors involved in the differentiation of thalamic neurons.

## Results

### *Sox2* and *Nr2f1* deletion in the developing visual thalamus causes deregulation of gene expression that precedes thalamic defects

To unravel the origin of the common phenotype observed in *Sox2* and *Nr2f1* thalamo-specific mouse mutants, we first stained the dLGN at postnatal day 0 (P0) (Fig. 1A) with anti-SOX2 and anti-NR2F1 antibodies and observed high co-expression of the two proteins in thalamic differentiated neurons (Fig. 1A). Then, to identify the gene regulatory network downstream of SOX2 and NR2F1 in the visual thalamus, we performed RNA-seq experiments on *ex vivo* dissected dLGN from *Sox2* or *Nr2f1* thalamic mutants (obtained via RORα-Cre deletion) and their control littermates at postnatal day 0 (P0), before the appearance of overt morphological impairments^7,9^ (Fig. 1B). Three independent pools of mutant and control dissected visual thalami for each mutant line were processed. We identified 2284 differentially expressed genes (DEG) with FDR < 0.01 following *Sox2* conditional inactivation, of which 1002 downregulated and 1282 upregulated (Fig. 1C; Table S1). In addition, with the same thresholds, we identified 1081 genes differentially expressed following *Nr2f1* conditional inactivation, of which 493 genes downregulated and 588 upregulated (Fig. 1C; Table S1). Notably, neither is *Nr2f1* dysregulated in *Sox2* mutant dLGN, nor is *Sox2* significantly dysregulated in the *Nr2f1* mutant, ruling out mutual regulation of the two genes (Table S1, see also ^9^). This was also confirmed by immunofluorescence showing no changes in the number and distribution of NR2F1-expressing cells in the absence of SOX2 (Fig. 1A), suggesting that the two proteins likely do not regulate each other, but control a common set of genes.

**Figure 1.**
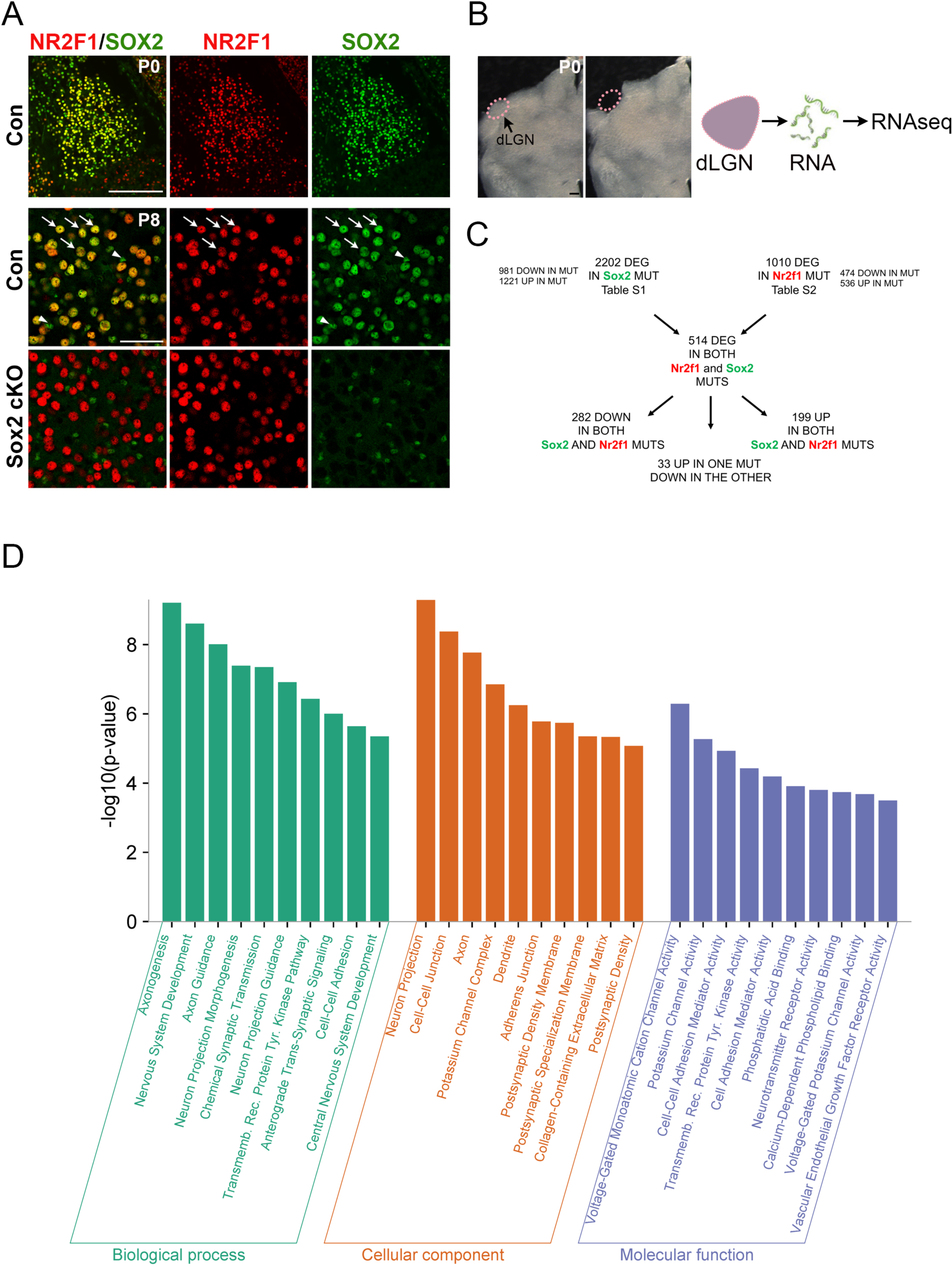
RNA-seq of the visual thalamus (dLGN) of *Sox2* and *Nr2f1* thalamic mutants identifies many genes differentially expressed in both mutants. **A**. dLGN immunofluorescence with antibodies recognizing SOX2 (green) and NR2F1 (red) at P0 (top; wild type) and P8 (bottom; wild type control, Con, and thalamic mutant, cKO). Note that Sox2 thalamic loss (*Sox2* cKO in the P8 panel) does not detectably affect NR2F1 expression. Scale bar top: 200um; scale bar bottom: 50um. **B**. Sections comprising the dLGN at P0 used for RNA-seq analyses, before (left) and after (right) dissection. Scale bar: 200um. **C**. Numbers of significantly dysregulated genes, identified in RNAseq experiments, in *Sox2* and *Nr2f1* mutants, and in both mutants are shown. **D.** Gene Ontology (GO) analysis of genes differentially expressed in both *Sox2* and *Nr2f1* mutants (DEG; the 514 genes in B) reveals enrichment in categories involved in neuronal development. The GO Biological Processes, Cellular Component and Molecular Function categories which are significantly enriched within the indicated mutants are shown.

### Many genes are co-regulated by SOX2 and NR2F1

We previously showed that important effectors of SOX2 function in NSC self-renewal and differentiation were among the most highly expressed and the most highly down-regulated genes in *Sox2*-mutated cells^13–15^. Importantly, genes dysregulated in *Sox2* mutant dLGN strikingly differed from those dysregulated in NSC^13^. We thus focused on the most down- or up-regulated genes in both thalamic mutants (Tables 1,2: Tables S2, S3). We first asked whether there are common genes deregulated in both *Sox2* and *Nr2f1* thalamic mutants. Fig. 1C shows that 514 genes significantly change their expression levels in both mutants, nearly all in the same direction (UP, or DOWN).

This figure by far exceeds the number of genes expected by pure chance (more than four times the expected value for a random overlap, and probability of having a similar overlap by chance < 10^-100^). Moreover, almost all the 50 genes in *Nr2f1* mutants with the most significant degree of downregulation are also significantly downregulated in *Sox2* mutants (44 out of 50); similarly, the majority of the 50 most downregulated genes in *Sox2* mutants are significantly downregulated also in *Nr2f1* mutants (30 out of 50) (Tables 1,2). Overall, these results clearly point to a SOX2 and NR2F1 co-regulated gene regulatory network in the early postnatal visual thalamus.

### Genes regulated by SOX2 and NR2F1 are enriched in functions related to neuronal differentiation and connectivity

To get insights into the collective functions of dysregulated genes, we performed Gene Ontology (GO) and functional enrichment analyses, focusing our attention on the genes significantly varying their expression in both *Sox2* and *Nr2f1* mutants (Fig. 1D). Results showed a striking enrichment in functions related to neuronal differentiation, and in particular to neuronal connectivity, activity and synaptic plasticity (Fig. 1D). Among the differentially expressed genes (DEG) represented in these categories, regulators in axon guidance (Efna5, EphA5, EphA7, Sema7A), specifically in retinal axon guidance (Nrp1, Alcam, EphB1), in glutamatergic synapses (Grid1, Cdh8), as well as transcription factors involved in visual development and axonogenesis (Sox5), could be identified.

Notably, among genes commonly downregulated in *Sox2* and *Nr2f1* mutants (Tables 1,2), ***Vgf*** deserved a particular attention, as it represented the most highly expressed, and one of the most strongly downregulated genes identified in both mutants (Tables 1,2). *Vgf* encodes a diffusible cytokine, transported along thalamo-cortical axons until the axon terminals. Its function is important for the development of cortical layer 4, onto which it acts instructively to maintain the appropriate numbers of layer 4 neurons, in particular within the somatosensory and visual cortical areas^16^. We thus looked at layer 4 development in thalamic *Sox2* mutants, using *in situ* hybridization for the layer 4 marker *RORβ*, the same marker previously used by ^16^ (Fig. 2A).

**Figure 2.**
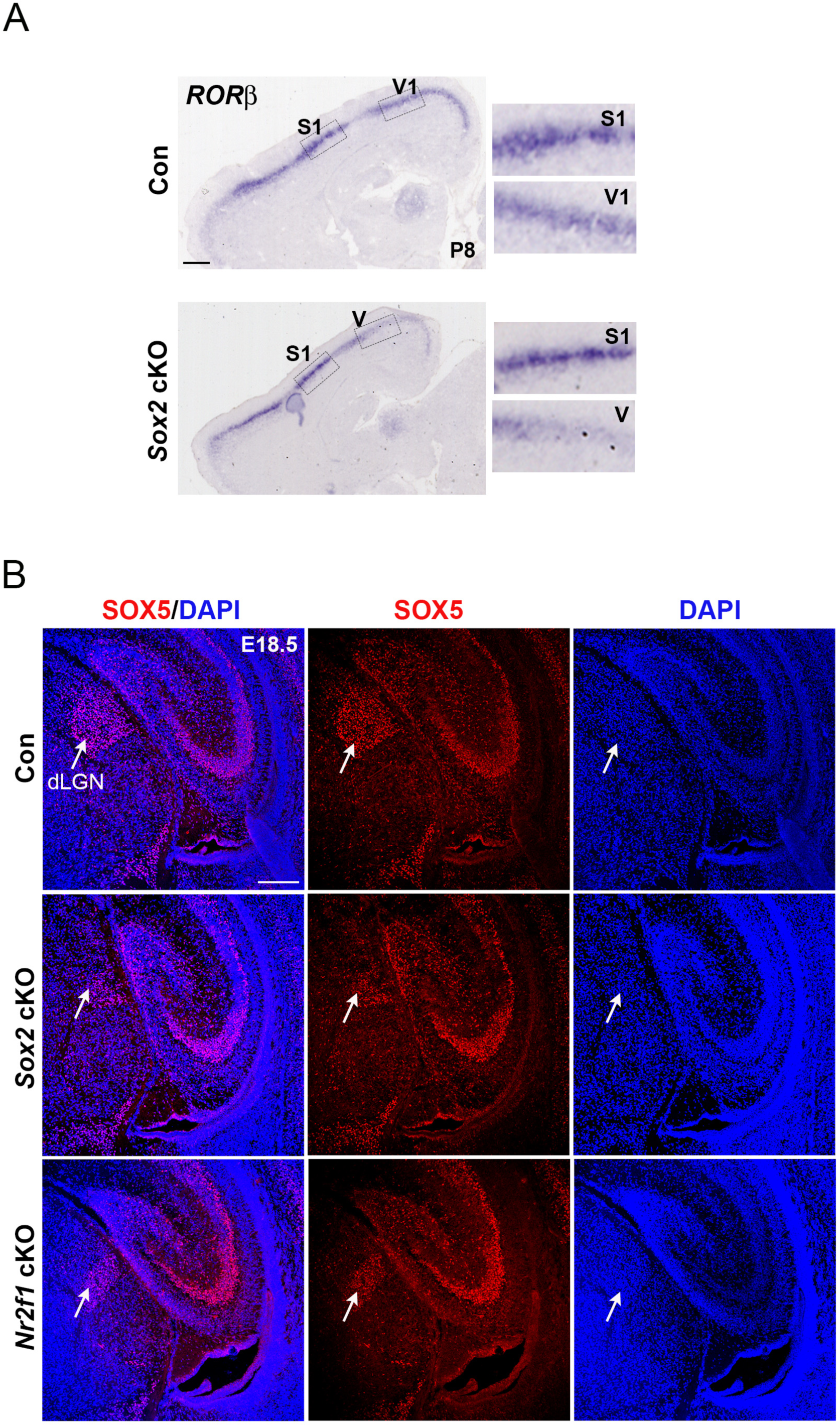
***Sox2* thalamic cKO affects the development of cortical layer 4, and thalamic SOX5 protein expression. A.** *In situ* hybridization with a *Rorβ* probe, marking layer 4, on sagittal brain sections of thalamic *Sox2* mutants (Sox2 cKO) versus controls carrying wild type *Sox2* (Con) at P8 shows a reduction in the region corresponding to the visual (V,V1) and somatosensory (S1) areas (enlarged details). The results shown are representative of n=3 mutant and n=3 control brains analysed. Scale bar: 600um. **B.** Immunofluorescence with antibodies recognizing SOX5 (red) on coronal sections of E18.5 brains of controls, *Sox2* cKO and *Nr2f1* cKO. DAPI (blue) marks nuclei. At this stage, the dLGN is only modestly reduced in the mutants^7,9^. Arrows point to SOX5-positive cells within the dLGN region. Note the reduction of the SOX5-positive area in the *Sox2* and *Nr2f1* mutant dLGN. SOX5 positivity is not altered in the adjacent hippocampal region in mutants. The results shown are representative of n=3 mutant and n=3 control brains analysed. Scale bar: 200um.

Indeed, we observed a reduction of the *RORβ* signal in the visual area (Fig. 2A), as observed in *Vgf*- knock-out mice^16^. This suggests that VGF may be a relevant downstream mediator of SOX2 thalamic function onto the development of cortical layer 4 in the visual area.

***Sox5***, encoding a transcription factor key to the development of cortical neuronal connectivity^17,18^, was also strongly downregulated in both mutants (Tables 1,2). Immunofluorescence studies with an anti-SOX5 antibody showed that the numbers of cells expressing SOX5 protein at high levels typical of wild-type cells were strongly diminished in both mutants (Fig. 2B).

### Deconvolution analysis identifies specific cell types affected by *Sox2* or *Nr2f1* deletion

A recent single-cell RNA-seq (scRNA-seq) analysis of the postnatal developing visual thalamus^11^, defined specific cell types by their distinct transcriptional identity. Specific gene expression patterns characterizing excitatory neurons, inhibitory neurons, oligodendrocytes, astrocytes, endothelial cells, pericytes and microglia could be defined, in the normal (wild type) situation. We thus performed a deconvolution analysis of our RNA-seq data, based on these single-cell transcriptional profiles at P5, the earliest time point analyzed by ^11^(Fig. 3), in order to estimate the abundance of each cell type in our samples. This showed a relevant reduction of the estimated fraction of glutamatergic neurons, in both *Sox2* and (to a lesser extent) *Nr2f1* mutant dLGNs (Fig. 3A,B).

**Figure 3.**
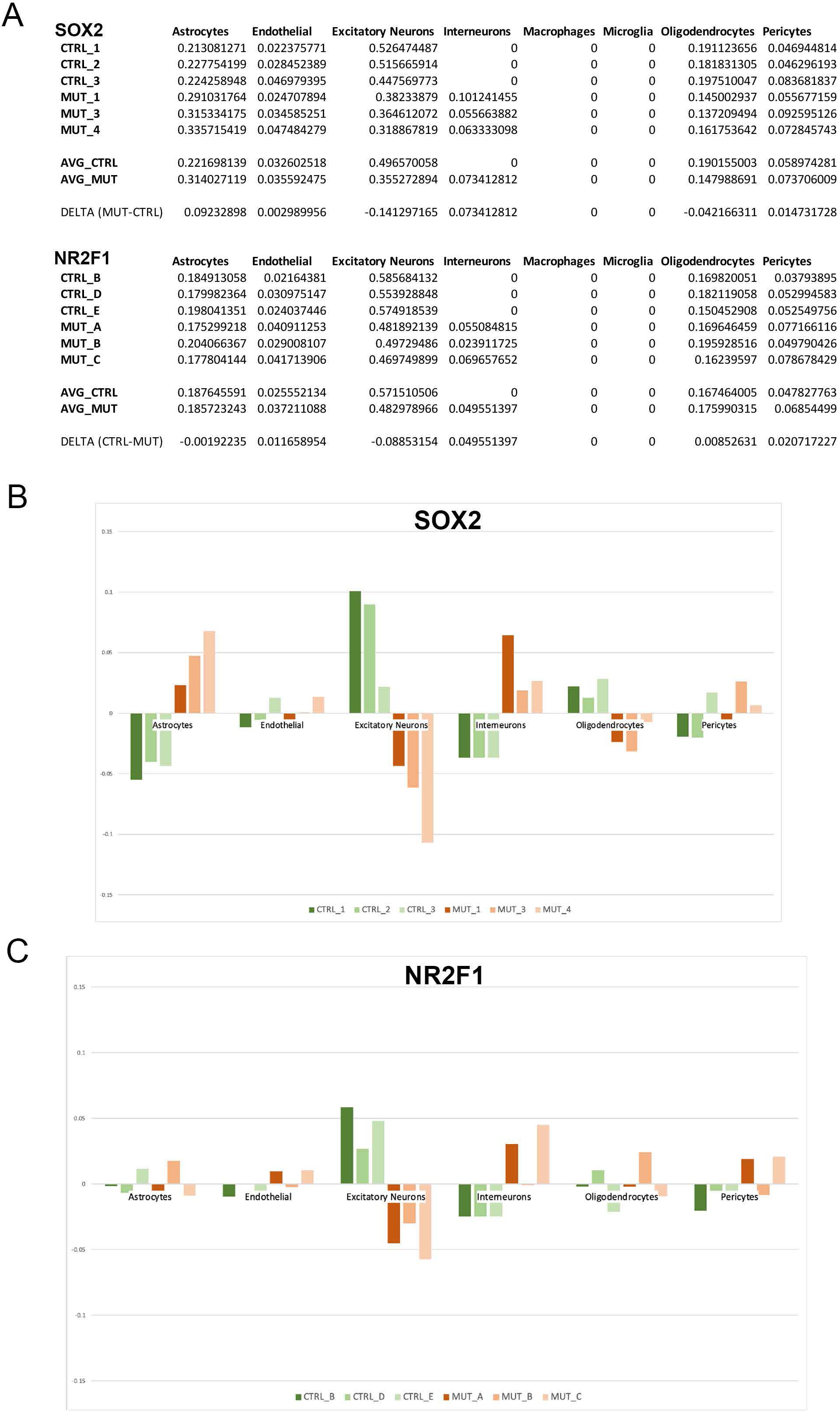
Deconvolution analysis documents the loss of projection neurons transcriptional identities in *Sox2* and *Nr2f1* mutants. The cell types indicated are those defined by their scRNA-seq transcriptional identity in ^11^. The data for all three mutant and control samples are shown numerically in (**A**) and graphically in (**B,C**).

Interneurons, pericytes and endothelial cells, conversely, were slightly increased, in both mutants. Oligodendrocytes were slightly reduced in *Sox2* mutants and increased in *Nr2f1* mutants. Of note, these changes were detected prior to overt abnormalities in the cell type composition of the mutant thalami, as previously shown by immunofluorescence and *in situ* hybridization only a week later, at P7 and/or P8^9^.

### CUT&RUN identifies SOX2 binding sites in the P0 visual thalamus

We set out determine whether the functionally relevant deregulated genes are direct targets of SOX2 by establishing the genome-wide binding profile of this transcription factor. To overcome the anticipated technical difficulties due to the scant cell number obtained by dissection of this structure, we pooled dLGNs from P0 wild-type newborns and subjected them to CUT&RUN targeting SOX2^12^ (Fig. 4A). Several thousand SOX2 peaks were identified in two independent experiments; their overlap further defined a group of 717 high-confidence, reproducible binding events (Fig. 4B). SOX2 peaks were found in promoter and intronic, as well as intergenic regions, consistent with the genomic binding pattern identified in other cellular systems ^13–15^(Fig. 4C). Two types of global analyses supported the dataset: a) Motif analysis revealed SOX2 as the top enriched motif, along with other SOX factors, likely due to motif similarities (Fig. 4D), increasing our confidence of the specificity of the identified peak regions. b) Gene ontology of peak-associated genes (by GREAT, ^19^) revealed enrichment for primarily neuronal biological processes (Fig. 4E), supporting involvement in the development of neurons and their connectivity.

**Figure 4.**
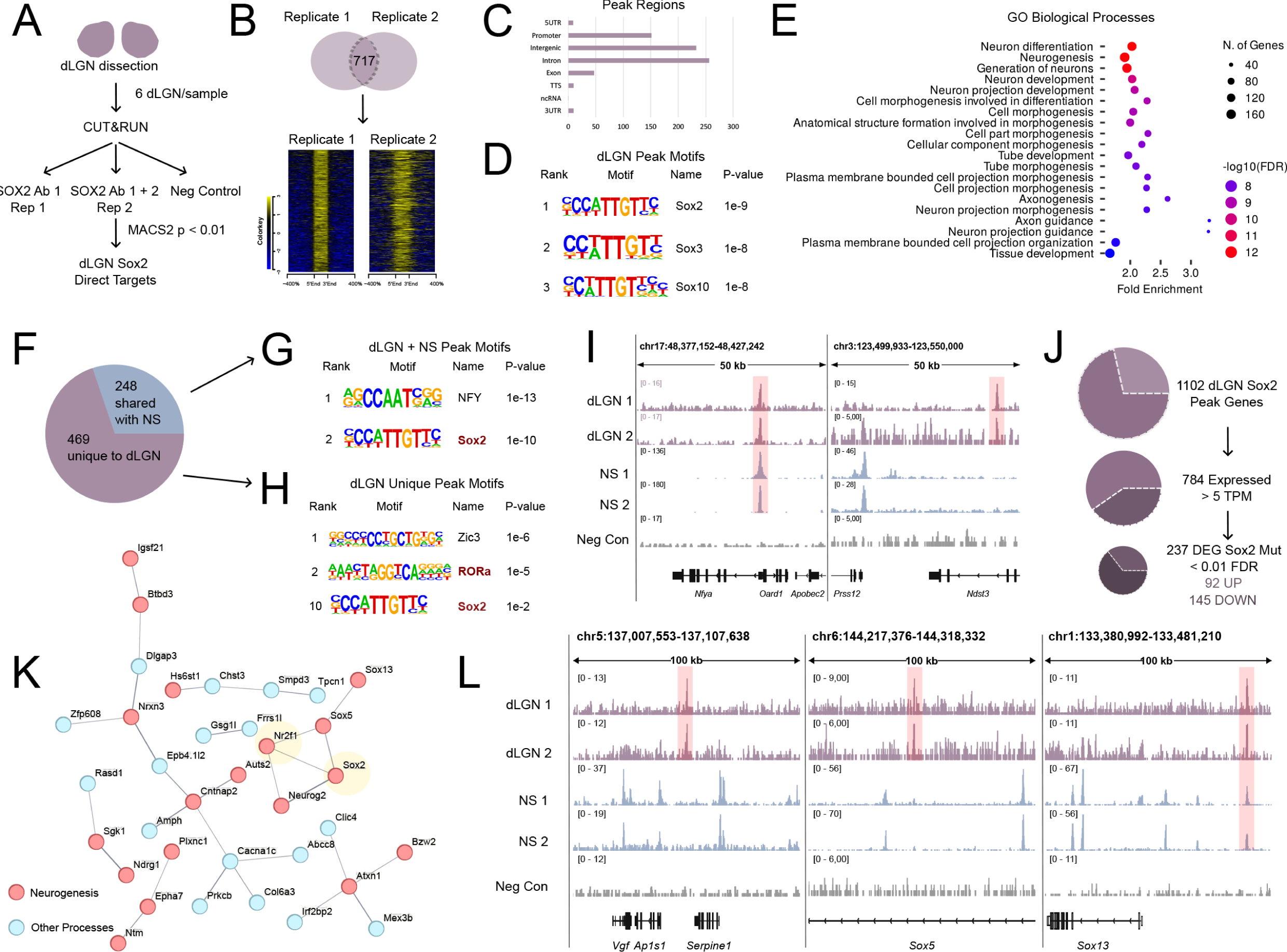
CUT&RUN of SOX2 binding in the P0 visual thalamus. A. Schematic depiction of the CUT&RUN experimental design. 2 independent biological replicates for SOX2 and an anti-HA negative control were performed from pools of 6 dorsal lateral geniculate nuclei (dLGN) from 3 brains. **B.** Venn diagram overlap and signal intensity plots of the two SOX2 replicates, showing enrichment over the control in all peak regions. **C.** Peak region annotation by HOMER, showing that SOX2 binds primarily promoter, intronic and intergenic regions in dLGN. **D.** HOMER known motifs for dLGN SOX2 peaks. The SOX2 motif is the most highly ranked, followed by other SOX factors. **E.** Gene ontology enrichment of biological processes for genes associated by GREAT to SOX2 dLGN peaks. Dot size shows number of peak associated genes, dot color represents -log10 FDR (FDR < 0.05), and the x axis represents fold enrichment. The top 20 terms are shown. Enrichment terms include neuron development related processes. **F.** Pie chart depicting the distribution of SOX2 dLGN peaks in unique peaks and those shared with neural stem cell (NS) SOX2 datasets. **G.** HOMER known motifs for dLGN and NS shared peaks. NFY and SOX2 are the top motifs. **H.** HOMER known motifs for dLGN unique peaks. Top motifs include ZIC3 and RORα. Sox2 is ranked 10^th^. **I.** CUT&RUN tracks as visualized in Integrative Genome Viewer (IGV), showing both dLGN and NS shared peaks (left) and dLGN unique peaks (right). **J.** Schematic depiction of CUT&RUN and RNA-seq overlap, showing genes that are transcribed ( > 5 TPM) and those that are differentially expressed (DEG) in *Sox2* mutant dLGN (FDR < 0.01). **K**. STRING map of interactions between SOX2, NR2F1, and the 79 SOX2 dLGN direct targets that are dysregulated in both *Sox2* and *Nr2f1* mutant mice. Confidence was set on default (0.4), disconnected nodes were removed, and line thickness represents confidence. Red nodes are those known to be involved in neurogenesis, while blue nodes are not included in the neurogenesis set and could represent novel genes important to the generation of neurons in the visual thalamus. **L.** CUT&RUN tracks showing SOX2 dLGN peaks near the important targets *Vgf* (left), *Sox5* (center, intronic region), and *Sox13* (right). While the peaks near *Vgf* and *Sox5* are unique to the dLGN datasets, the *Sox13* peak is also bound by SOX2 in NS.

We compared the genome-wide physical occupancy of SOX2 in the dLGN to our previously identified binding of this factor in brain-derived neural stem cells (NSC) performed both with CUT&RUN and with ChIP-seq^14,15^. Among the 717 high-confidence targets, 248 were shared with NSC (Fig. 4F). This allowed us to define a subset of 469 dLGN specific SOX2 targets (Fig. 4F), which likely underlay a different role of SOX2 in this differentiated cell population. Of note, while the shared targets almost only displayed motif enrichment for SOX factors (Fig. 4G), the dLGN- specific SOX2 peak regions contained an enrichment for the RORα consensus sequence, highly similar to the NR2F1 binding site – a motif that is not enriched in the neural stem cell subset (Fig. 4H). Examples of peaks shared with the neural stem cells and unique to the dLGN can be seen in Fig. 4I,L; note the presence of peaks in the proximity of the *Vgf* (Fig. 4I) and intronically within *Sox5* (Fig. 4L) regions, which are among the most strongly downregulated genes in *Sox2* mutant (see above and Table 1), suggesting their direct regulation by SOX2.

**Table 1.** Genes most down-regulated in *Sox2*-mutant visual thalamus.

| Gene | Av. Exp.<br>**Sox2 WT**<br>TPM | Av. Exp.<br>**Sox2 MUT**<br>TPM | Av. Exp. **Sox2 WT**/<br>Av. Exp. **Sox2 MUT** | log fold<br>change | DEG in<br>**Nr2f1 MUT** |
| --- | --- | --- | --- | --- | --- |
| 9030625G05Rik | 8.1 | 1.05 | 7.69 | -2.895780452 | **YES** |
| Hdc | 21.82 | 2.97 | 7.35 | -2.813501459 |  |
| Fam83f | 9.84 | 1.6 | 6.15 | -2.550812898 | YES |
| Ptpn22 | 4.52 | 0.84 | 5.4 | -2.371974499 | YES |
| Tuba8 | 9.12 | 1.75 | 5.2 | -2.313158562 |  |
| Hs6st2 | 109.4 | 22.54 | 4.85 | -2.230994808 |  |
| Pcdhac1* | 14.92 | 3.04 | 4.9 | -2.224814845 | **YES** |
| Drd5 | 10.05 | 2.23 | 4.51 | -2.107744688 | YES |
| Flt3 | 11.74 | 2.72 | 4.31 | -2.044844104 | YES |
| Wnt9b | 50.83 | 12.19 | 4.17 | -1.997868077 | **YES** |
| Hs3st1 | 121.89 | 29.48 | 4.13 | -1.986194868 |  |
| Rora | 62.54 | 16.24 | 3.85 | -1.955983475 |  |
| Nexn | 7.58 | 1.92 | 3.95 | -1.927547838 | **YES** |
| Neurog2 | 12.08 | 3.05 | 3.96 | -1.925343754 | **YES** |
| Hs6st3* | 48.32 | 12.18 | 3.97 | -1.92121348 | **YES** |
| Vgf | 882.88 | 231.32 | 3.82 | -1.873781013 | **YES** |
| Muc15 | 6.6 | 1.72 | 3.84 | -1.870164907 | **YES** |
| Sp9* | 269.34 | 78.82 | 3.42 | -1.710807965 |  |
| Lgi2 | 35.57 | 10.41 | 3.42 | -1.707749283 |  |
| Slc18a2 | 218.62 | 64.43 | 3.39 | -1.701630392 |  |
| Col9a1 | 9.37 | 3 | 3.13 | -1.698584742 |  |
| Cpne9 | 43.55 | 13.27 | 3.28 | -1.655753749 | YES |
| Efna5 | 37.94 | 11.68 | 3.25 | -1.638069788 |  |
| Cd47 | 129.27 | 41.23 | 3.14 | -1.586910507 | YES |
| Tmem132d | 20.26 | 6.61 | 3.07 | -1.551311996 | **YES** |
| Galnt14 | 34.04 | 11.15 | 3.05 | -1.548701903 |  |
| Drd1 | 4.73 | 1.55 | 3.04 | -1.543540066 |  |
| 2900005J15Rik | 6.47 | 2.13 | 3.04 | -1.539759249 |  |
| Chrm3 | 16.91 | 5.67 | 2.98 | -1.512314896 | **YES** |
| Adgrl2 | 88.59 | 29.81 | 2.97 | -1.510982218 | YES |
| Tmem132b* | 70.73 | 23.74 | 2.98 | -1.510191854 | **YES** |
| Edaradd | 9.05 | 3.05 | 2.97 | -1.507508032 |  |
| Frrs1l | 71.66 | 24.07 | 2.98 | -1.506989596 | YES |
| Zmat4* | 25.97 | 8.78 | 2.96 | -1.50054383 | **YES** |
| Kcnn1 | 45.79 | 15.68 | 2.92 | -1.484704706 |  |
| Tmem163 | 142.8 | 48.96 | 2.92 | -1.484221758 | YES |
| Ephx4 | 9.52 | 3.3 | 2.88 | -1.469356885 | YES |
| Trhde | 12.26 | 4.25 | 2.89 | -1.461311822 |  |
| Sox13 | 86.24 | 30.48 | 2.83 | -1.440847204 | YES |
| Cdca7 | 51.7 | 18.3 | 2.83 | -1.43834782 |  |
| Dmrtb1* | 111.05 | 39.94 | 2.78 | -1.416756694 | **YES** |
| Cpne4 | 34.3 | 12.39 | 2.77 | -1.406373382 | **YES** |
| Slc6a4 | 58.61 | 21.26 | 2.76 | -1.402243263 | **YES** |
| Fzd8 | 17.56 | 6.36 | 2.76 | -1.399855218 |  |
| Sox5 | 25.07 | 9.1 | 2.75 | -1.397136566 | YES |
| Dkk4 | 6.33 | 2.33 | 2.71 | -1.374842774 | **YES** |
| Il22 | 7.01 | 2.6 | 2.69 | -1.372466032 |  |
| Camk4 | 88.09 | 32.64 | 2.7 | -1.367842536 | YES |
| Gm15417 | 17.04 | 6.48 | 2.63 | -1.346050718 | YES |
| Dusp5 | 8.72 | 3.31 | 2.63 | -1.335628473 |  |

**Table 2.**
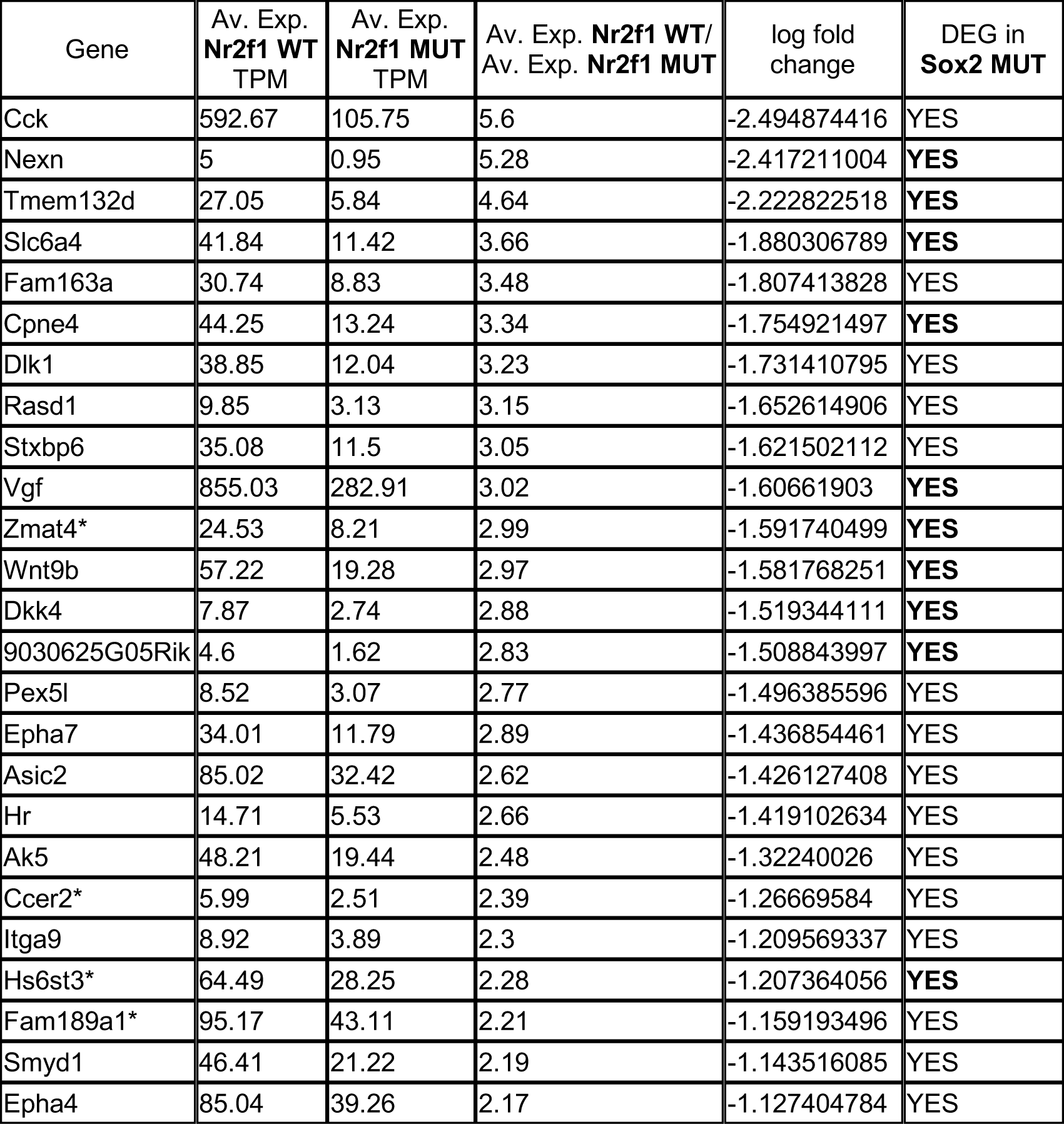
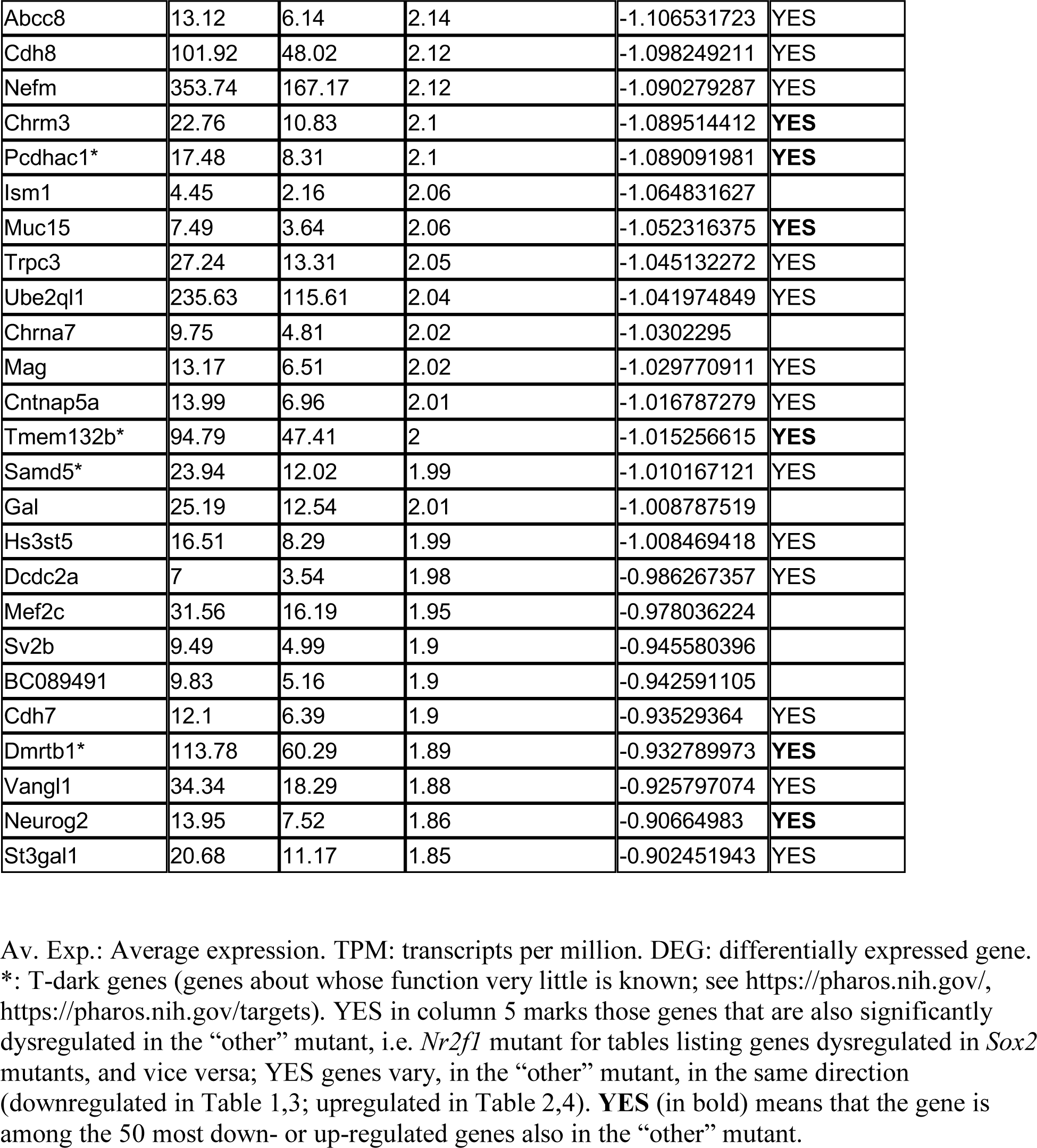
Genes most down-regulated in *Nr2f1*-mutant visual thalamus

GREAT annotated the 717 binding events to a total of 1102 genes (Fig. 4J). This allowed us to overlap our CUT&RUN with the genes identified to be expressed or differentially expressed when Sox2 is inactivated (RNA-sequencing data, Fig. 1C; Table S1). We found that 784 of the 1102 peak-associated genes were expressed in dLGNs at P0 (TPM > 5). Of these, 247 (145 down, 92 up) were dysregulated in SOX2 mutant dLGNs (< 0.01 FDR) (Fig. 4J), providing robust evidence of SOX2 direct regulation in differentiated neurons.

To understand whether these genes could also be regulated by NR2F1, we overlapped these differentially expressed SOX2 targets with the genes dysregulated in *Nr2f1* mutant dLGNs. This led to a core signature of 79 SOX2 and NR2F1 coregulated genes, which display known literature-based physical and functional interactions (Fig. 4K, STRING ^20^ diagram of core interacting SOX2/NR2F1 coregulated genes, disconnected nodes removed). Of note, this list includes several genes that are among the top 100 up- or down- regulated upon SOX2 deletion in the dLGN, including *Dmrtb1*, *Sgk1*, *Cnr2*, *Sox5*, *Gsg1l*, *Frrs1l*, *Sox13*, *Flt3*, *Neurog2*, *Hk3* (Fig. 4L).

## Discussion

The conditional deletion of *Sox2*, and of *Nr2f1*, from the developing visual thalamus leads to important, significantly overlapping defects in the thalamus itself, as well as in the thalamus- connected visual cortex^7,9^. We show that more than 1000 genes are dysregulated in each of the two mutants. Among them, an important subset of genes downregulated in both *Sox2* and *Nr2f1* mutants points to the existence of a shared network of genes essential for the proper development of the dLGN and its connections. In principle, the genes downregulated in the *Sox2*- and the *Nr2f1*- mutant dLGNs should explain three types of phenotypic defects observed in mutant mice^7,9^: (i) the reduction in the number of neurons (glutamatergic) in the mutant visual thalami (at postnatal day 7); (ii) the alterations in terms of amount and distribution of thalamo-cortical and cortico-thalamic connectivity; (iii) the alterations in the retino-geniculate connections.

Indeed, gene ontology (GO) analysis of the genes dysregulated in both mutants reveals a highly significant enrichment of categories related to functions that develop abnormally in the mutants. These include, as the most significantly enriched categories, biological processes such as axonogenesis, neuron projection morphogenesis, chemical synapsis transmission, axon development, retinal ganglion cell axon guidance (Fig. 2D), and related cellular components (Fig. 2D). It is likely that, collectively, genes forming these categories underlie, by their altered regulation, the above-mentioned defects.

### Potential key contributors of SOX2- and NR2F1-dependent vision development

As shown in Tables 1,2, a large proportion of genes highly expressed in WT cells, and highly down-regulated in *Sox2* mutants, are also down-regulated in *Nr2f1* mutants; the reciprocal is also true. In contrast, among the genes up-regulated in *Sox2* mutants, just a few are upregulated also in *Nr2f1* mutants (Tables S2, S3). This suggests that the functional association of SOX2 and NR2F1 is mostly involved in positive regulation (activation) of gene transcription.

### Vgf

Among the genes at the top of the list, *Vgf* is the most highly expressed and among the most strongly downregulated ones, among those downregulated in both *Sox2* and *Nr2f1* mutants (Table 1). *Vgf* encodes a signaling molecule, transported along thalamo-cortical axons until the axon terminals in the L4 layer of the cortex, where it acts instructively to maintain the appropriate neuron numbers within the visual and somatosensory cortical areas ^16^. The role of VGF in the layer was initially demonstrated by ablating the thalamo-cortical neuronal connections, thus depriving the cortex of sufficient VGF. Further, the knock-out of *Vgf* strongly reduced layer 4 in both the somatosensory and visual cortices. This phenotype could then be rescued to normality by transgenic expression of VGF in the developing cortex ^16^. We observe (Fig. 2A) that *RORβ*, marking cortical layer 4, is moderately reduced in the visual cortex of *Sox2* mutants, mirroring, at least to an extent, what is found in *Vgf* mutants ^16^. Knowing that VGF expression is strongly reduced in the mutant dLGN (Table 1,2), and that axonal connections reaching the visual cortex are also reduced in both mutants ^7,9^, and in agreement with the abnormalities in the primary (V1) and higher order (V^HO^) visual cortical areas in both thalamic mutants ^7,9^, we hypothesize that the observed cortical layer defects of the *Sox2* mutant (Fig. 2A) might be the consequence of a reduced amount of VGF reaching the axon terminals in the visual cortex. Thus, VGF might be a relevant mediator of the function of SOX2 during area- and layer-specific cortical development. Furthermore, *Sox2* is also down-regulated in the somatosensory VP (Ventro-Posterior) thalamic nucleus in *Sox2* thalamo- specific conditional mutants, leading to moderate abnormalities of the somatosensory cortical area^9^. We thus looked at the somatosensory cortex layer 4 and observed a *RORβ* reduction (less marked than in the visual area) also in this area (Fig. 2A), in agreement with the above hypothesis.

### Sox5

SOX5, a transcription factor, is significantly downregulated in both mutants at the mRNA and protein level (Table 1,2; Fig. 2B). In humans, the *SOX5* gene is associated to “Optic nerve hypoplasia bilateral, autosomal dominant” (OMIM #165550). As shown in *Sox5-*null mice ^18^ ^17^ this gene controls important aspects of neuronal connectivity, such as the development of cortico- fugal neurons, including cortical neurons projecting to thalamic neurons. Misrouting of subplate and layer 6 cortico-thalamic axons to the hypothalamus is also observed ^17^. These data suggest the possibility that the reduction of SOX5 in the dLGN of *Sox2* and *Nr2f1* mutants affects axonogenesis.

### Hs6st2, Hs6st3

The Heparan sulphate 6-O-sulfotransferase enzyme modifies the sulfation status of heparan sulphate proteoglycans (HSPG) ^21^, extracellular matrix proteins, with covalently linked polysaccharidic chains, which are polymerized by specific enzymes and further modified by sulfation to obtain ample structural and functional diversity ^22^. HSPG interact with a variety of proteins, implicated in cell proliferation and differentiation, adhesion, migration, and other processes ^21^. It is intriguing that three of the genes (*Hs6st1*, *2* and *3*) encoding isoforms of the enzyme are downregulated in *Sox2* and/or *Nr2f1* mutant dLGNs. Axon guidance or axon extension defects abnormalities, and eye defects, have been reported in association with mutation of *Hs6st* genes^21–25^. We hypothesize that SOX2- and NR2F1-dependent expression of HS6ST enzymes, acting on the sulfation status of HSPGs, might play a role in the development of axonal connectivity of thalamic neurons.

### Rorα

*Rorα*, also named *Nr1f1*, is one of the most downregulated genes in *Sox2* mutants, although not in *Nr2f1* mutants (Tables 1,2); yet, it may have a special significance also in the perspective of the present study, focusing on a SOX2 and NR2F1 co-regulated gene expression network. *Rorα* encodes a close homolog of NR2F1, and its product NR1F1 binds to a DNA sequence closely related to that of NR2F1, with an identical core binding site (https://www.genecards.org/cgi-bin/carddisp.pl?gene=RORA; https://www.genecards.org/cgi-bin/carddisp.pl?gene=NR2F1). *Rorα* spontaneous (staggered mouse) or engineered mutations cause ataxia and cerebellar neurodegeneration, with synaptic arrangement and immature morphology of cerebellar neurons (Purkinje) ^26,27^*. RORα* defects are connected in humans with “Intellectual development disorder with or without epilepsy, or cerebellar ataxia, OMIM #600825. If the activity of RORα on some genes partially overlaps with that of NR2F1, it is possible that, in the *Sox2* mutant, the reduced activity of RORα leads to down-regulation of a set of genes closely related to those activated by NR2F1, explaining some common phenotypic defects observed in *Sox2* and *Nr2f1* mutants.

### Mechanisms of dysregulation of gene expression in mutant thalamus

The observed dysregulation of several genes in the mutant thalami could be due to variations in the abundance of specific cell types, as well as to variations in gene expression levels within specific cell types normally expressing SOX2 and NR2F1. They may also be due to indirect effects, whereby decreased expression of genes downregulated following *Sox2* or *Nr2f1* knockout affects the expression of other genes not directly targeted by SOX2 or NR2F1 themselves. Evidence for direct effects of *Sox2* or *Nr2f1* deficiency is provided by CUT&RUN , which identified several hundreds of chromatin sites bound by SOX2 in replicate experiments (Fig. 4).

The variation in cell numbers in the mutants is indeed important. In fact, in previous work, we detected a reduction in the number of neurons by P7 following *Sox2* thalamic deletion ^9^. In the present work, we detect a specific reduction in the fraction of cells bearing a transcriptional program characteristic of projection neurons, already at P0 (Fig. 3, deconvolution). This reduction is likely to be rooted in gene expression abnormalities within glutamatergic neurons themselves, which normally express SOX2 and NR2F1, and not in a decreased number of cells. Accordingly, several of our downregulated genes (e.g. *Vgf*, *Sox5*, *Rorα*) are reported to be prevalently expressed in neurons in published datasets of scRNA-seq analyses ^28^. On the other hand, the large relative increase in GABAergic interneurons (that do not express SOX2 in the thalamus)^9^ might be due to indirect effects, such as a change in the fate of part of the mutant neurons, or an increased interneuron colonization of the mutant thalamus, or increased survival of the interneurons in the mutant thalamus.

### dLGN-specific SOX2 binding sites differ from those shared with neural stem cells

The comparison of SOX2 binding sites detected by CUT&RUN in the visual thalamus (present paper) with those previously observed in neural stem cells by both CUT&RUN and ChIPseq ^13–15^ allowed us to compare the most represented SOX2 binding sequence motifs specific to the thalamus with those specific to NSC. SOX2 binding sites are among the most represented sites in both datasets, as expected (Fig. 4D, 4G, 4H). Remarkably, however, the thalamus-specific SOX2- binding sites showed, among the most represented, the binding site for RORα, also called NR1F1. As discussed above, NR1F1 and NR2F1, highly homologous, recognize similar DNA sequences with a core reported to be identical (https://www.genecards.org/cgi-bin/carddisp.pl?gene=RORA; https://www.genecards.org/cgi-bin/carddisp.pl?gene=NR2F1). This suggests that the molecular basis for the sharing of co-regulated target genes between SOX2 and NR2F1 might lie in the co-binding of the two factors to regulatory DNA sequences.

Overall, our work identifies a common transcriptional program driven by SOX2 and NR2F1 in the visual thalamus, highlighting a small subset of genes commonly downregulated in both mutants, and potentially important for explaining abnormalities of retina-thalamus-cortex connections. This paves the way to the precise identification, by functional transgenic studies, of genes mediating the common Sox2 and Nr2f1 functions in the visual thalamus. It will also open the way to defining the molecular mechanisms mediating SOX2 and NR2F1 interactions in transcriptional regulation.

## Materials and Methods

### Mouse strains

Sox2 Mutant mice were obtained by crossing Sox2Flox (Favaro et al., 2009) with Rorα-Cre (Chou et al., 2013) mouse lines.

Nr2f1 mutant mice were generated by crossing COUP-TF1Flox (Armentano et al., 2007) with Rorα- Cre (Chou et al., 2013) mouse lines (note: COUP-TF1 is the old name of Nr2f1).

Genotyping was performed with the following primers (Chou et al., 2013; Mercurio et al. 2019): Rorα-Cre IRES Forward: 5’AGGAATGCAAGGTCTGTTGAAT 3’; Rorα-Cre IRES Reverse: 5’ TTTTTCAAAGGAAAACCACGTC 3’; Sox2 Flox Forward: 5’AAGGTACTGGGAAGGGACATTT 3’; Sox2 Flox Reverse: 5’AGGCTGAGTCGGGTCAATTA 3’; COUP-TF1 Flox Forward 5’-CTGCTGTAGGAATCCTGTCTC-3’; COUP-TF1 Flox Reverse: 5’- AATCCTCCTCGGTGAGAGTGG-3’ and 5’– AAGCAATTTGGCTTCCCCTGG-3’.

The day of vaginal plug was defined as embryonic day 0 (E0) and the day of birth as postnatal day 0 (P0).

We note that Rorα is the gene to the 3’ of which the Cre gene has been inserted (as IRES-Cre knocked-in into the 3’UTR of the gene), in the Cre transgene driving Sox2 and Nr2f1 deletion in our thalamic mutants. In principle, *a priori*, the Cre insertion into the Rorα locus per se might have lowered Nr1f1-Rora expression. However, we also note that the significant (about 4 times) reduction of expression is observed in Sox2 mutants, but not Nr2f1 mutants, where expression levels are unchanged with respect to control. We thus think it is unlikely that Cre insertion plays a role in the expression reduction of Rora in Sox2 mutants.

### Brain extraction and dissection for RNA-sequencing

Brains from Sox2 thalamic mutants and controls, and Nr2f1 thalamic mutants and controls, at postnatal day 0 (P0) were removed from the skulls in ice-cold PBS1x, coronally embedded in low melt agarose 4% in PBS1x and kept on ice. Brains were then sectioned in ice-cold and sterile PBS1x with a vibratome (Leica VT1000s). 200µm brain sections were collected in ice cold PBS1x. Sections including the dLGN were identified under a stereoscopic microscope and the dLGN was quickly dissected with sterile chirurgic scalpels on a glass slide. dLGNs were present usually in two sequential sections. Excised dLGNs were collected with a pipette, collected in sterile tubes, snap-frozen in liquid nitrogen and stored at -80°C until RNA extraction. All the sections were imaged before and after dissection of dLGNs.

### RNA extraction and RNA sequencing

RNA from snap-frozen dLGNs was extracted with the RNeasy Micro Kit (QIAGEN) with some precautions. Each dissected dLGN weighed about 0,3 mg. dLGNs samples were taken out of the - 80°C and left few seconds on the bench before tissue homogenization. Each dLGN sample was homogenized in 350 µl of RLT buffer containing β-mercaptoethanol (β-ME) as indicated by the manufacturer (10 µl of β-ME per 1 ml of RLT buffer). Tissue was homogenized by sequential passages through needles of descending diameter. Precisely we used a 1cc syringe and three different sterile needles: 18G, 22G and 26G. First, the tissue was passed 10 times through the larger diameter needle (18G), then 10 times through the intermediate diameter needle (22G) and finally 10-12 times through the thinnest one (26G), in order to correctly lysate and disrupt the tissue. Subsequently, the homogenized sample was vortexed for 30 seconds and centrifuged for 3 minutes at 13200 rpm to pellet the debris. RNA was extracted from the homogenized supernatant as indicated by the manufacturer’s instructions.

RNA sequencing was performed on three independent samples for both mutant and control dLGN. Each sample was composed of dLGNs from three animals of the same genotype pooled together. Genotypes were: for mutants, Sox2flox/flox, or Nr2f1flox/flox, plus RORalpha-Cre transgene; for controls, we used littermates of the respective mutants (Sox2 or Nr2f1 mutants), carrying two intact copies of Sox2 (Sox2flox/flox or Sox2flox/+), or Nr2f1 (Nr2f1flox/flox or Nr2f1flox/+), and no Cre transgene. For each sample sequenced we obtained at least 150 ng of high quality total RNA (RIN ≥ 8), and thus 150 ng were used for library preparation. Library preparation was performed with Nugen Universal + mRNAseq kit, followed by sequencing in a HiSeq 4000 (Illumina), paired-reads 2x 75 bp. The KEGG 2019 Mouse database was used to analyse RNA-sequencing data.

The Gene Ontology analysis reported in Fig. 1 was done using Enrichr (https://maayanlab.cloud/Enrichr/0).

### RNA-seq raw data analysis and deconvolution analysis

Sequence reads were mapped with STAR^29^ against the mouse RefSeq transcriptome, version April 2019, retrieved from the UCSC Genome Browser Database^30^. Read counts and subsequent normalized transcripts per million (TPM) were computed with RSEM (rsem-calculate-expression)31.

Differential expression analysis was performed with edgeR ^32^. Initial read counts were normalized by trimmed mean of M values (TMM), with default parameters. Differentially expressed genes were identified by the quasi-likelihood (QL) F-test of edgeR (glmQLFfit and glmQLFtest functions, with default parameters). We selected as differentially expressed all genes with an adjusted p-value (FDR) < 0.01.

Bulk RNA-Seq deconvolution analysis was performed with MuSiC ^33^, with as input raw count tables for bulk RNA-Seq and raw count tables at P5 for single cell RNA-Seq samples. Single cell clusters and cell type annotations were retrieved from ^11^.

For Tables 1-4, DEG were initially selected having expression levels higher than 4 TPM (at least in wild type, for downregulated genes; at least in mutant, for upregulated genes). Average expression values were then calculated from triplicate samples in Table S1. The top 50 most down- or up- regulated genes (with greatest log-fold change in mutant versus wild type) are shown for both mutants.

### Immunohistochemistry

To detect SOX2 and NR2F1, slides were washed 2X for 10 minutes in PBS1X and then unmasked in Na Citrate 0.1M-Citric acid 0.1M pH6 solution for 2 minutes. Sections were washed in PBS1X for 10 minutes and incubated 1 h with pre-blocking solution (Sheep serum 5%, Tween-20 0,3% in PBS1X) at room temperature. Sections were then incubated over night at 4°C in blocking solution (Sheep serum 1%, Tween-20 0,1% in PBS1X) containing primary antibodies. The following primary antibodies were used: anti-SOX2 diluted 1:500 (R&D AB2018, mouse) and anti-NR2F1 diluted 1:1000 (Abcam Ab181137, rabbit). Sections were then washed 2X for 10 minutes in PBS1X and incubated for 1h at room temperature with blocking solution containing the following fluorescent secondary antibodies: anti-mouse IgG Alexa Fluor 488 and anti-rabbit IgG Alexa Fluor 594 (Thermo Fisher) diluted 1:500. Slides were then washed 2X for 10 minutes in PBS1X and mounted.

To detect SOX5, slides were treated as above, but without the unmasking step. The anti-SOX5^34^ antibody (a gift from A. Morales) was diluted 1:500, and the anti-rabbit Alexa Fluor 594 (Thermo Fisher) was diluted 1:1000.

### *In situ* hybridization

*In situ* hybridization was performed as in (Mercurio et al. 2019). Briefly, brains at P7 brains were dissected and fixed overnight in paraformaldehyde (PFA) 4% in PBS (Posphate Buffered Saline) 1X. The fixed tissue was cryoprotected in a series of sucrose solutions (15%, 30%) in PBS 1X and then embedded in OCT (Killik, Bio-Optica) and stored at -80°C. Brains were sectioned (20 µm) with a cryostat, placed on a slide (Super Frost Plus 09-OPLUS, Menzel) and stored at -80°C. Slides were then defrosted, fixed in formaldehyde 4% in PBS for 10 minutes (min), washed 3 times for 5 min in PBS 1X, incubated for 10 min in acetylation solution (for 200 ml: 2.66 ml triethanolamine, 0.32 ml HCl 37%, 0.5 ml acetic anhydride 98%) with constant stirring and then washed 3 times for 5 min in PBS1X. Slides were placed in a humid chamber and covered with prehybridization solution (50% formamide, 5X SSC, 0.25 mg/ml tRNA, 5X Denhardt’s, 0.5 µg/ml salmon sperm) for at least 2 hours (h) and then incubated in hybridization solution (fresh prehybridization solution containing the digoxygenin (DIG)-labelled RNA probe of interest) overnight at 65°C. Slides were washed 5 min in 5X SSC, incubated 2 times in 0.2X SSC for 30 min at 65°C, washed 5 min in 0.2X SSC at room temperature and then 5 min in Maleic Acid Buffer (MAB, 100 mM maleic acid, 150 mM NaCl pH 7.5). The slides were incubated in blocking solution (10% sheep serum, 2% blocking reagent (Roche), 0.3% Tween-20 in MAB) for at least 1 h at room temperature, then covered with fresh blocking solution containing anti-DIG antibody Roche © 1:2000 and finally placed overnight at 4°C. Slides were washed in MAB 3 times for 5 min, in NTMT solution (100 mM NaCl, 100 mM Tris-HCl pH 9.5, 50 mM MgCl2, 0.1% Tween-20) 2 times for 10 min and then placed in a humid chamber, covered with BM Purple (Roche), incubated at 37°C until desired staining was obtained (1-6 h), washed in water for 5 min, air dried and mounted with Eukitt (Sigma). A DIG-labelled *RORβ* probe was used^35^.

### CUT&RUN

CUT&RUN was performed as described in ^36^. dLGNs were dissected (as described above in Brain extraction and dissection for RNA-sequencing), suspended in 1.5 ml of cold CUT&RUN wash buffer (HEPES pH 7.5 [20 mM], NaCl [150 mM], Spermidine [0.5 mM], Roche Complete Protease Inhibitor EDTA-Free (Cat. # COEDTAFRO, Roche) and then manually dissociated by gentle pipetting. Cells were collected by centrifugation at 600g for 3 minutes and the supernatant carefully removed, and then washed 2 more times by resuspension in 1 ml wash buffer. 4 µl of ConA bead slurry (prepared in binding buffer as described), were added per pair of dLGNs and the cells and beads were incubated for 10 min on a rotator. While the rest of the dissections proceeded, bead bound cells were kept in storage buffer (wash buffer with EDTA [2 mM]). 3 brains (6 dLGNs) were collected per sample. Once all dLGNs were collected, the beads were collected on the magnet and resuspended in 150 µl antibody buffer (wash buffer with EDTA [2 mM] and 0.025 % digitonin). Antibodies were added at 1:100 dilution and incubation proceeded ON at 4 °C. Antibodies used included anti-SOX2 (ABIN2855074, antibodies online), anti-SOX2 (ABIN2855073, antibodies online), and anti-HA (05- 902R, Merck). One SOX2 sample was with only ABIN2855074, the other was with both SOX2 antibodies together. The two SOX2 biological replicates were performed independently from different litters of mice. The next day samples were washed twice with dig-wash (wash buffer with 0.025% digitonin) and then resuspended in 150 µl dig-wash containing pA-MNase (New England Biolabs 700 ng/ml, received as a gift from Steven Henikoff) at 1:200 dilution and rotated for 1 hr. Samples were washed twice and then resuspended in 100 µl dig-wash and equilibrated in ice. 2 µl 100 mM CaCl2 was added, and digestion proceeded for 30 min in ice. 100 µl 2X STOP buffer (NaCl [340 mM], EDTA [20 mM], EGTA [4 mM], digitonin [0.05%], RNase A [100 µg/ml], glycogen [50 µl/ml]) was added to stop the digestion reaction and incubated at 37 °C for 30 min. Samples were pelleted at 16000 g for 5 min and placed on the magnet rack. The supernatant was harvested, and beads were discarded. DNA purification was performed with phenol chloroform. Library preparation was performed using the KAPA HyperPrep Kit (Cat. #KK8504, KAPA Biosystems) according to manufacturer’s instructions, using KAPA DUI adapters at 0.15 µM. Libraries were sequenced with the Illumina NextSeq 550 using the High-Output 75 cycles kit v2.5 (Cat. #20024906, Illumina), 36 base pair pair-end.

### CUT&RUN Data Analysis

Reads were trimmed to remove adapters, artifacts and repeats of poly [AT]36, [C]36 and [G]36 with bbmap bbduk ^37^ (version 38.18). Alignment was performed to the mm10 mouse genome with bowtie ^38^ (version 1.0.0) with the options -v 1 -m 1 -X 500. Samtools ^39^ (version 1.11) was used for deduplication and to remove and incorrectly paired reads. Bedtools ^40^ (version 2.30.0) was used to remove reads mapped to the CUT&RUN mm10 Suspect List ^41^ from bam files. Peaks were called for each replicate using MACS2 ^42^ with the options -f BAMPE --keep-dup all -p 1e-2 -SPMR -bdg against the anti-HA negative control. Output narrowPeak files were overlapped using Bedtools intersect, keeping only reproducible peak regions called in both biological replicates. Signal intensity plots were created using ngsplot (https://doi.org/10.1186/1471-2164-15-284, version 2.63) with options -N 4 -GO none -SC global, plotting signal intensity of the replicates compared to the anti-HA control. Motif analysis was done using HOMER ^39^ (version 4.11) findMotifsGenome to find motifs in the mm10 genome using -size given, and peak region annotation was done with HOMER annotatePeaks on default settings. Peak set gene annotation was done using GREAT ^19^ (version 4.0.4) with default parameters, and gene names were used to compare CUT&RUN results with the RNA-seq analysis. Gene ontology was performed for GO biological processes using ShinyGO ^43^ (https://doi.org/10.1093/bioinformatics/btz931), results were cutoff at FDR of 0.05 and the top 20 results were graphed. For neural stem cell CUT&RUN data, datasets were downloaded from ^44^ and processed as described above for the dLGN datasets. The two peak sets were overlapped with Bedtools intersect. STRING ^45^ was used to graph protein-protein interactions between the set of 79 genes that were found to be dLGN SOX2 targets coregulated by SOX2 and NR2F1, disconnected nodes were removed. Bedgraphs were visualized in IGV ^46^.

### Accession to genomic data

RNA-seq data are accessible through Gene Expression Omnibus (GEO) Accession Number GSE233131. CUT&RUN data are accessible through ArrayExpress Accession Number E-MTAB- 13004.

## Acknowledgements

We thank Dr. D. O’Leary for providing the RORα-Cre mouse, Dr. Carolina Frassoni and her laboratory for contributing to the immunofluorescence experiments in Fig. 1A, and Dr. Aixa Morales for supplying the anti-SOX5 antibody. Work in the Nicolis laboratory is supported by EU ERANET-NEURON grant Brain4Sight and by Fondazione Telethon – Fondazione Cariplo Alliance GJC21176 grant to S.K.N. Work in the Cantù lab is supported by Cancerfonden (CAN 2018/542 and 21 1572 Pj), the Swedish Research Council, Vetenskapsrådet (2021–03075), Linköping University support and LiU-Cancer. C.C. is a Wallenberg Molecular Medicine (WCMM) fellow and receives generous financial support from the Knut and Alice Wallenberg Foundation. The computations and data handling for CUT&RUN analyses were enabled by resources provided by the National Supercomputer Centre (NSC), funded by Linköping University. Peter Münger at the National Supercomputer Centre is acknowledged for assistance concerning technical and implementational aspects in making the codes run on the Sigma resource.

