## Supplementary material for "SOX2 and NR2F1 coordinate the gene expression program of the early postnatal visual thalamus": Tables S1, S2, S3: Tables S2 S3.pdf

**Table S2**
**Genes most up-regulated in Sox2-mutant visual thalamus**

| Gene | Av. Exp. **Sox2 WT** TPM | Av. Exp. **Sox2 MUT** TPM | Av. Exp. **Sox2 WT**/ Av. Exp. **Sox2 MUT** | log fold change | DEG. in **Nr2f1 MUT** |
|---|---|---|---|---|---|
| Dclk3 | 3.25 | 13.07 | 0.25 | 2.069804975 | |
| Nts | 2.14 | 7.83 | 0.27 | 1.937609382 | |
| Ntng2 | 3.54 | 11.65 | 0.3 | 1.756206937 | |
| Gbx2 | 18.98 | 58.25 | 0.33 | 1.68474955 | YES |
| Nrgn | 21.16 | 63.12 | 0.34 | 1.644634092 | |
| Trem2 | 6.83 | 19.64 | 0.35 | 1.585346573 | |
| Lhx6 | 1.9 | 5.23 | 0.36 | 1.532534867 | |
| Agt | 19.03 | 51.7 | 0.37 | 1.502400404 | |
| Lyz2 | 6.1 | 16.58 | 0.37 | 1.485172692 | |
| Baiap3 | 3.82 | 10.09 | 0.38 | 1.464322031 | YES |
| Sstr2 | 1.6 | 4.1 | 0.39 | 1.464037449 | |
| P2ry12 | 5.94 | 15.69 | 0.38 | 1.459736983 | |
| Doc2b | 4.27 | 11.15 | 0.38 | 1.455343586 | |
| Plscr5* | 7.97 | 20.56 | 0.39 | 1.423373905 | **YES** |
| Ccl3 | 1.71 | 4.39 | 0.39 | 1.414103031 | |
| Hist1h1c | 5.87 | 15.01 | 0.39 | 1.403302332 | |
| Kirrel3 | 24.54 | 62.01 | 0.4 | 1.402286818 | YES |
| Lpl | 3.72 | 9.3 | 0.4 | 1.387522058 | |
| Aif1 | 3.19 | 8.07 | 0.4 | 1.384584124 | |
| Ramp3 | 7.08 | 17.67 | 0.4 | 1.384117839 | YES |
| Evi2a | 3.18 | 7.94 | 0.4 | 1.379864659 | |
| Crabp2 | 18.68 | 46 | 0.41 | 1.35623076 | |
| Rph3al | 1.8 | 4.41 | 0.41 | 1.355173978 | |
| Ntf3 | 2.35 | 5.76 | 0.41 | 1.351913409 | |
| Drd2 | 4.02 | 9.8 | 0.41 | 1.347035815 | |
| C1qb | 27.74 | 67.81 | 0.41 | 1.345230759 | YES |
| Kcnab1 | 1.72 | 4.18 | 0.41 | 1.34021734 | |
| Nxpe4 | 2.14 | 5.2 | 0.41 | 1.337769469 | |
| Tyrobp | 21.1 | 51.42 | 0.41 | 1.330841028 | |
| Vsnl1 | 55.46 | 133.2 | 0.42 | 1.326367214 | |
| Syt17 | 2.98 | 7.14 | 0.42 | 1.319653648 | |
| Hck | 1.81 | 4.32 | 0.42 | 1.319231384 | |
| Fcrls | 14.58 | 34.76 | 0.42 | 1.312364771 | YES |
| Adgre1 | 2.94 | 7 | 0.42 | 1.309497369 | |
| Sst | 37.25 | 88.18 | 0.42 | 1.303432488 | |
| Elmod1 | 7.53 | 17.68 | 0.43 | 1.299201494 | |
| Eln | 19.92 | 47.48 | 0.42 | 1.296634367 | **YES** |
| Calb1 | 4.2 | 9.8 | 0.43 | 1.290495697 | |
| Lhx1os | 2.78 | 6.58 | 0.42 | 1.289104521 | |

| | | | | |
|---|---|---|---|---|
| Gfap | 56.27 | 131.77 | 0.43 | 1.288362339 |
| Pcsk2 | 18.96 | 44.05 | 0.43 | 1.282187768 |
| C1qa | 27.03 | 62.78 | 0.43 | 1.270652334 |
| Renbp | 5.55 | 12.84 | 0.43 | 1.260929494 |
| Sla | 2.86 | 6.51 | 0.44 | 1.258201969 |
| Slc17a7 | 5.42 | 12.36 | 0.44 | 1.254317242 |
| Synpr | 1.96 | 4.45 | 0.44 | 1.251098849 |
| Grm1 | 3.59 | 8.19 | 0.44 | 1.25062982 |
| Calb2 | 130.53 | 297.43 | 0.44 | 1.249230203 |
| Crabp1 | 4.86 | 11.13 | 0.44 | 1.247895487 |
| Kcnj5 | 1.84 | 4.15 | 0.44 | 1.240846025 |

**Table S3**
**Genes most up-regulated in Nr2f1-mutant visual thalamus**

| Gene | Av. Exp. **Nr2f1 WT** TPM | Av. Exp. **Nr2f1 MUT** TPM | Av. Exp. **Nr2f1 WT**/ Av. Exp. **Nr2f1 MUT** | log fold change | DEG in **Sox2 MUT** |
|---|---|---|---|---|---|
| 4930426D05Rik | 1.73 | 14.95 | 0.12 | 3.07842899 | |
| Klhl14 | 3.56 | 24.99 | 0.14 | 2.787445111 | |
| Adamts16 | 1.79 | 11.3 | 0.16 | 2.629609081 | |
| Acta2 | 2.21 | 11.46 | 0.19 | 2.318237041 | |
| Crispld2 | 1.48 | 5.78 | 0.26 | 1.939943191 | |
| Aifm3 | 2.69 | 9.73 | 0.28 | 1.765572256 | |
| Ccl21a | 4.62 | 15.41 | 0.3 | 1.721195858 | YES |
| Gjb6 | 1.52 | 4.97 | 0.31 | 1.677811435 | |
| Siah3 | 5.09 | 16.56 | 0.31 | 1.675559409 | |
| Slc22a6 | 3.53 | 11.01 | 0.32 | 1.607191342 | |
| Syndig1* | 4.42 | 13.65 | 0.32 | 1.601930695 | YES |
| Oxtr | 3.16 | 9.43 | 0.33 | 1.568033413 | YES |
| 6430573F11Rik | 1.34 | 4.06 | 0.33 | 1.557353524 | |
| Spsb1 | 5.8 | 17.28 | 0.34 | 1.550556523 | |
| Ebf1 | 24.4 | 71.48 | 0.34 | 1.530242289 | |
| Brinp2 | 24.66 | 70 | 0.35 | 1.482398431 | |
| 1700086L19Rik | 4.69 | 13.15 | 0.36 | 1.481287226 | |
| Plscr5* | 4.16 | 11.73 | 0.35 | 1.480475238 | **YES** |
| Tekt1 | 3.03 | 8.53 | 0.35 | 1.479901939 | |
| Eln | 15.91 | 43.96 | 0.36 | 1.436271118 | **YES** |
| Slc35f4* | 2.44 | 6.72 | 0.36 | 1.433830311 | |
| Dbpht2 | 2.08 | 5.69 | 0.36 | 1.424776402 | |
| Gpr139 | 2.67 | 7.28 | 0.37 | 1.416859524 | |
| Tinagl1 | 1.87 | 5.11 | 0.37 | 1.413944585 | |
| Angptl2 | 2.75 | 7.27 | 0.38 | 1.375583156 | |
| Mrgprf | 1.88 | 4.9 | 0.38 | 1.353667852 | |

| | | | | | |
|---|---|---|---|---|---|
| Ppp1r32 | 2.36 | 6.12 | 0.39 | 1.35151314 | |
| Tac1 | 2.53 | 6.3 | 0.4 | 1.331242164 | |
| Edn3 | 3.41 | 8.61 | 0.4 | 1.303340784 | |
| Procr | 2.29 | 5.71 | 0.4 | 1.296105068 | |
| Dera | 3.38 | 8.37 | 0.4 | 1.287440734 | YES |
| Greb1 | 1.87 | 4.56 | 0.41 | 1.282017302 | |
| Mpped2 | 11.81 | 28.82 | 0.41 | 1.275560135 | |
| Dact2 | 1.75 | 4.31 | 0.41 | 1.275337329 | YES |
| Trpm8 | 1.91 | 4.68 | 0.41 | 1.275256445 | |
| Mfap4 | 17.19 | 42.42 | 0.41 | 1.273352892 | |
| Clec3b | 9.27 | 22.84 | 0.41 | 1.269238615 | |
| Gpr182 | 2.65 | 6.47 | 0.41 | 1.264855514 | YES |
| Sidt1 | 8.49 | 20.74 | 0.41 | 1.262250086 | |
| Mrc1 | 6.33 | 15.39 | 0.41 | 1.254676366 | |
| Ngef | 9.08 | 22.01 | 0.41 | 1.2523175 | |
| Kctd16 | 3.58 | 8.65 | 0.41 | 1.24984524 | |
| Phf24 | 23.36 | 56.11 | 0.42 | 1.246183299 | |
| Amn | 4.37 | 10.59 | 0.41 | 1.245180361 | |
| Slc6a13 | 19.46 | 46.57 | 0.42 | 1.230500284 | |
| Gyltl1b | 1.85 | 4.29 | 0.43 | 1.217651861 | |
| Ptgds | 85.22 | 198.26 | 0.43 | 1.184876577 | YES |
| Twist2 | 2.28 | 5.25 | 0.44 | 1.177026019 | |
| Igsf9 | 10.92 | 24.91 | 0.44 | 1.173048534 | YES |
| Arhgap10 | 6.55 | 14.58 | 0.45 | 1.135771598 | |

Av. Exp.: Average expression. TPM: transcripts per million. DEG: differentially expressed gene. *: T-dark genes (genes about whose function very little is known; see https://pharos.nih.gov/, https://pharos.nih.gov/targets). YES in column 5 marks those genes that are also significantly dysregulated in the "other" mutant, i.e. Nr2f1 mutant for tables listing genes dysregulated in Sox2 mutants, and vice versa; YES genes vary, in the "other" mutant, in the same direction (downregulated in Table 1,3; upregulated in Table 2,4). **YES** (in bold) means that the gene is among the 50 most down- or up-regulated genes also in the "other" mutant.
